# Conditional deletion of LRRC8A in the brain protects against stroke damage independently of effect on swelling-activated glutamate release

**DOI:** 10.1101/2022.12.13.520314

**Authors:** Mustafa Balkaya, Preeti Dohare, Sophie Chen, Alexandra L. Schober, Antonio M. Fidaleo, Julia W. Nalwalk, Rajan Sah, Alexander A. Mongin

## Abstract

The ubiquitous volume-regulated anion channels (VRACs), which are composed of LRRC8 proteins, facilitate cell volume homeostasis, and contribute to many other physiological processes. Prior studies demonstrated that treatment with non-specific VRAC blockers, or brain-specific deletion of the essential VRAC subunit LRRC8A, are highly protective in rodent stroke. In this work, we tested the widely accepted idea that harmful effects of VRACs in the brain are mediated by pathological release of the excitatory transmitter glutamate. We used two molecular genetic strategies to ablate LRRC8A expression in either brain astrocytes only (inducible deletion of *Lrrc8a*^flox/flox^ with *Aldh1l1*^CreERT2^) or the majority of brain cells (neurons, astrocytes, and oligodendrocytes with *Nestin*^Cre^). To produce stroke, genetically modified mice were subjected to a 40-minute occlusion of the middle cerebral artery. The inducible deletion of astrocytic LRRC8A yielded no histological or behavioral protection. In contrast, the brain-wide LRRC8A knockout reduced ischemic infarction by ~50% in both heterozygotes (Het) and the full *Lrrc8a* knockout (KO) as compared to the control *Lrrc8a*^flox/+^ genotype. However, despite identical brain damage, Het and KO mice dramatically differed in their VRAC activities. Het mice had full swelling-activated glutamate release, while KO animals showed its virtual absence. These new findings refute the notion that VRAC-mediated glutamate release plays significant role in ischemic brain injury.

## INTRODUCTION

Stroke is defined as a loss of brain functions caused by an interruption of cerebral blood flow, either due to vessel occlusion (ischemic stroke) or rupture (hemorrhagic stroke) (Sacco et al., 2013;Campbell et al., 2019). The high prevalence, rapid onset, and severity of tissue injury in stroke make this disease the second leading cause of death and the third leading source of disability worldwide (Campbell et al., 2019;Feigin et al., 2022). Despite the large burden on public health, the currently approved acute stroke therapies are limited to thrombolytic agents, most notably tissue plasminogen activator (tPA or Alteplase) and its genetically modified form Tenecteplase, or mechanical retrieval of clots (Campbell et al., 2019). Despite many promising preclinical studies, clinical trials have failed to confirm efficacy of neuroprotective agents in stroke patients (Ginsberg, 2008;Paul and Candelario-Jalil, 2021). Therefore, there is a pressing need for a better understanding of stroke pathology and the discovery of new therapeutic targets.

One prospective molecular target in stroke that gained attention over the past two decades are the volume-regulated anion channels (VRACs). VRACs are the ubiquitous chloride channels in vertebrates, which are activated in response to cellular swelling and play a canonical role in cell volume regulation (Strange et al., 1996;Okada, 1997;Nilius et al., 1997). Opening of VRACs initiates efflux of cytosolic anions, indirectly facilitating the loss of intracellular K^+^ and osmotically obligated water (Hoffmann et al., 2009;Pedersen et al., 2016;Jentsch, 2016). In addition to inorganic anions, such as Cl^-^ and bicarbonate, VRACs are permeable to various small organic molecules which carry either a negative charge, such as glutamate, aspartate, lactate, and pyruvate, or are net-neutral, e.g., taurine, sorbitol, glutamine, and GABA (Strange et al., 1996;Okada, 1997;Nilius et al., 1997). Since many amino acid neurotransmitters can permeate VRACs, in the brain their opening can result in hyperexcitability and, in extreme cases, lead to tissue injury (Kimelberg and Mongin, 1998;Kimelberg, 2005;Mongin, 2016).

The idea of VRAC involvement in stroke pathology stems from the observations that brain ischemia triggers robust cellular swelling and that swollen cells release large quantities of glutamate (Kimelberg, 2005;Mongin, 2007) This hypothesis has been tested using pharmacological tools, such as the estrogen receptor antagonist tamoxifen, which at micromolar levels potently inhibits VRAC, and the more selective DCPIB. In rodent stroke models, tamoxifen and DCPIB decrease the intraischemic buildup of glutamate and aspartate (Phillis et al., 1998;Seki et al., 1999;Feustel et al., 2004), and potently reduce infarction volumes and neurological deficits (Kimelberg et al., 2000;Kimelberg et al., 2003;Mehta et al., 2003;Feng et al., 2004). Yet, due to the off-target effects of both agents, the uncertainty of VRAC contribution to ischemic pathology remains unresolved. Recently, the field was revolutionized by the discovery that VRACs are comprised of proteins belonging to the Leucine-Rich Repeat-Containing family 8 (LRCC8) (Qiu et al., 2014; Voss et al., 2014). There are five LRRC8 proteins (LRRC8A-E). LRRC8A is essential for VRAC conductance but it must be combined with other LRRC8 homologues in order to form fully functional heterohexameric channel (Qiu et al., 2014;Voss et al., 2014;Deneka et al., 2018;Kasuya et al., 2018;Kefauver et al., 2018;Nakamura et al., 2020). It is now possible to use molecular genetics to definitively test biological roles for the LRRC8A subunit and VRACs.

Our present work follows two recent publications probing the role of LRRC8A in ischemic brain injury. The first study has found that astrocyte specific LRRC8A knockout moderately reduces brain damage in a murine model of stroke (Yang et al., 2019). A subsequent report revealed more robust protection in mice carrying the brain wide LRRC8A deletion (Zhou et al., 2020). These discoveries have strengthened the hypothesis that VRAC is an important player in stroke pathology. Yet, their conclusions are based on a correlation between LRRC8A expression and stroke outcomes rather than the direct testing LRRC8A involvement in pathological glutamate release. Another caveat is that conditional knockouts can lead to developmental adaptations. Indeed, the brain wide LRRC8A deletion produces reactive astrogliosis, gives rise to spontaneous seizures, and results in 100% animal mortality withing the first eight weeks of postnatal development (Zhou et al., 2020;Wilson et al., 2021). Therefore, in the present work, to avoid developmental compensation we tested the impact of LRRC8A deletion on stroke outcomes using an inducible deletion of LRRC8A in astrocytes. Because astrocytic ablation was not neuroprotective, we extended our efforts to the brain wide LRRC8A knockout. Surprisingly, our work reveals the lack of correlation between swelling-activated glutamate releases and stroke protection.

## RESULTS

### Validation of *Aldh1l1^CreERT2^-driven* excision of *Lrrc8a* in the CNS astrocytes

To produce inducible, astrocyte-specific deletion of LRRC8A, we utilized commercially available *Aldh1l1*^CreERT2^ mice, which was produced and validated by Khakh laboratory (Srinivasan et al., 2016). Aldehyde dehydrogenase 1 family member L1 (ALDH1L1) is a folate metabolism enzyme, whose expression in in the adult brain is nearly exclusive to astrocytes (Cahoy et al., 2008). Our breeding strategy yielded the *Aldh1L1*^CreERT2/+^; *Lrrc8a*^fl/fl^ genotype, hereafter referred to as inducible astrocytic LRRC8A knockout (iaLRRC8A KO) and *Lrrc8a*^fl/fl^ littermates (Figure 1A). Mice were treated with tamoxifen to induce Cre expression or corn oil to provide vehicle treatment control. Altogether, we analyzed four genotype/treatment groups with only one of them carrying the deletion of LRRC8A in astrocytes (Figure 1A).

**Figure 1:**
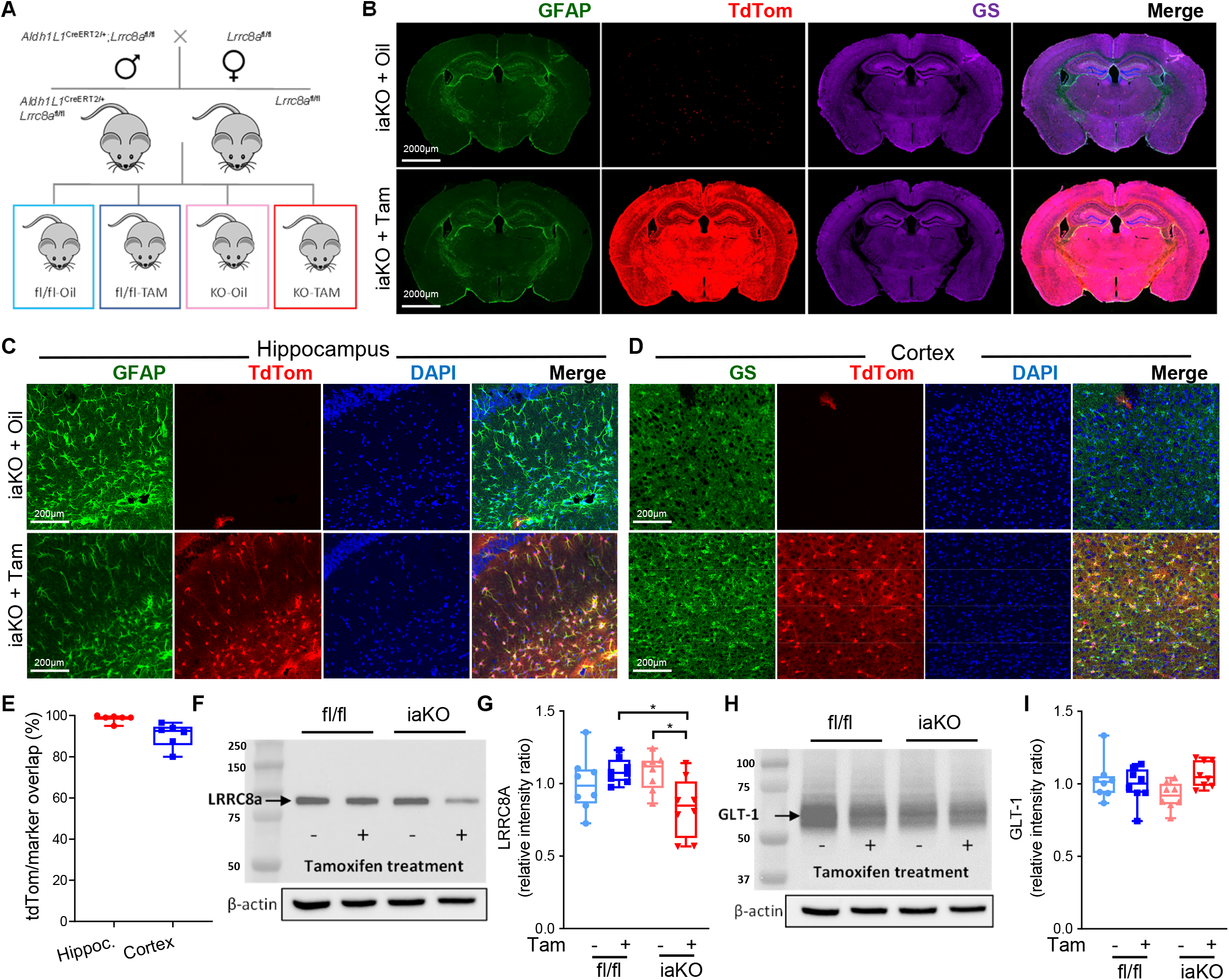
Validation of *Aldh1l1*^CreERT2^ driven LRRC8A deletion in mouse astrocytes *in vivo*. **(A)** Breeding and treatment strategy for the astrocyte-specific inducible deletion of LRRC8A using *Aldh1l1*^Cre^-targeted excision of *Lrrc8a*^fl/fl^. The *Aldh1l1*^CreERT2^;*Lrrc8a*^fl/fl^ mice (iaKO) and *Lrrc8a*^fl/fl^ (fl/fl) controls were treated with either tamoxifen (Tam, activates CreERT2) or vehicle (Oil) to generate four experimental groups. **(B)** Representative immunohistochemistry images of whole-brain sections from 12-week old iaKO mice showing expression of the Cre reporter tdTomato (red) and the astrocyte markers, GFAP (green) and glutamine synthetase (GS, purple). **(C)** Representative high magnification images (200×) visualizing the co-expression of tdTomato (red) and GFAP (green) in the hippocampus. **(D)** Representative high magnification images (200×) visualizing the coexpression of tdTomato (red) and GS (green) in the cortex. **(E)** Quantification of the colocalization of the immunoreactivity (IR) for the Cre reporter tdTomato and the astrocytic markers GFAP (in hippocampus) and GS (Cortex). Data are the mean values ± SD in six brains. **(F)** Representative western blot image of LRRC8A expression in the whole brain lysates from four genotype/treatment groups. **(G)** Quantification of LRRC8A expression. The box plot of individual IR values, normalized to β-actin loading and averaged within a set. n=8/group. *p<0.05, iaKO+Tam vs. fl/fl+Tam and iaKo+oil; two-way ANOVA with Sidak’s multiple comparisons test; ^##^p<0.01, genotype-treatment interaction. **(H)** Representative western blot image of GLT1 expression in four genotype/treatment groups. **(I)** Quantification of GLT1 expression. The box plot of individual IR values, normalized to β-actin loading and averaged within a set.

To validate effective LRRC8A deletion we performed two types of experiments. First, we utilized the inbred tdTomato reporter to visualize Cre activity in the brain. As seen in Figure 1B, tamoxifen treatment of iaLRRC8A KO mice produced robust tdTomato expression throughout the brain. In contrast, no tdTomato fluorescence was observed in the iaLRRC8A KO mice treated with corn oil, except for a few positive cells reflecting spontaneous Cre activity (Figure 1B). The *Lrrc8a*^fl/fl^ controls had no tdTomato fluorescence (data not shown). Higher magnification images demonstrated that tdTomato fluorescence overlapped with the astrocytic markers GFAP in hippocampus (Figure 1C) and glutamine synthetase in cortical regions (Figure 1D). We used glutamine synthetase in cortex because of low GFAP expression levels (see Figure 1B and review (Khakh and Sofroniew, 2015)). Extensive quantification in six brains demonstrated Cre induction in >98% hippocampal astrocytes and >90% cortical astrocytes (Figure 1E).

To substantiate LRRC8A protein deletion, we next performed semi-quantitative western blot analyses in whole-brain lysates. We found ~20% reduction of LRRC8A expression in the tamoxifen-treated iaLRRC8A KO mice, as compared to oil-treated iaLRRC8A KO group and tamoxifen-treated *Lrrc8a*^fl/fl^ controls (p<0.05, Figure 1F, G). Previously, we found that the brain-wide deletion of LRRC8A reduces astrocytic expression of the glutamate transporter GLT-1 (Wilson et al., 2021). Therefore, we additionally explored if there are changes in GLT-1 immunoreactivity in iaLRRC8A KO. No change in GLT-1 expression was found in any of tested groups (Figure 1H, I).

Next, to assess the extent of LRRC8A loss in astrocytes, we performed studies in primary astrocyte cultures prepared from neonatal iaLRRC8A KO brains. Astrocyte cultures were treated with either 2 μM hydroxy-tamoxifen (4-OHT) or vehicle (0.05% DMSO) for 4-14 days and analyzed for the expression levels of LRRC8A and induction of the Cre reporter, tdTomato. tdTomato reporter was expressed as early as 24 h, labeled >70% of cultured cells by days 3-4, and reached saturated intensity by days 6-8 (Figure 2A). Western blot analysis identified robust downregulation of LRRC8A protein expression, with a surprising lag of approximately 10 days (Figure 2B, C). The extent of LRRC8A deletion varied among cell preparations from nearly full (representative blot in Figure 2B) to partial, but on average reached ~80% by days 12 and 14 (Figure 2C). The temporal mismatch between changes in LRRC8A and tdTomato expression may indicate the different accessibility of these two genes for Cre.

**Figure 2:**
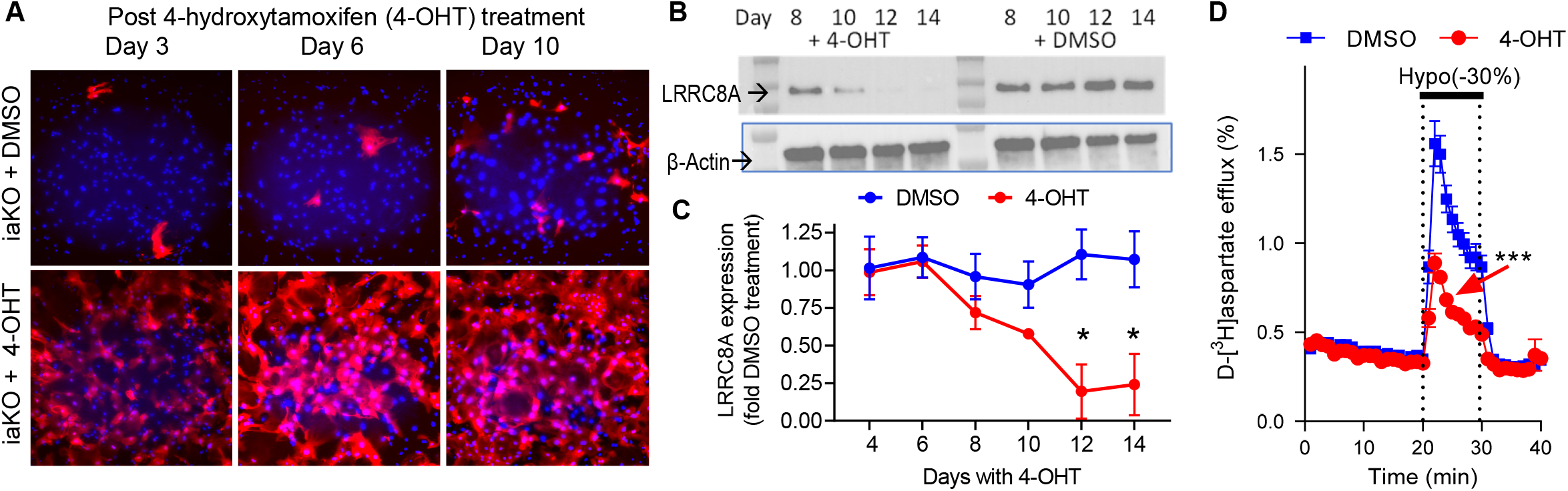
*Aldh1l1*^CreERT2^ driven reductions in LRRC8A expression and VRAC activity in mouse iaLRRC8A KO astrocytes *in vitro*. **(A)** Representative images of the Cre reporter tdTomato expression in the *Aldh1l1*^CreERT2^;*Lrrc8a*^fl/fl^ (iaKO) astrocyte cell cultures treated with either 2 μM 4-hydroxytamoxifen (4-OHT) or vehicle control (DMSO). Images show overlay of tdTomato (red) and Hoechst (blue) staining at days 3, 6 and 10 after initiation of treatment. **(B)** Representative western blot of LRRC8A expression in 4-OHT of vehicle (DMSO) treated astrocytic cultures at days 8 to 14 after initiation of treatment. **(C)** Quantification of LRRC8 expression. Data are the mean IR values of individual immunoreactivity ± SEM normalized to β-actin loading controls and to average values. n=3/group. p<0.05, 4-OHT vs DMSO at D12 and D14. Repeated measures ANOVA with Sidak’s post hoc comparison. **(D)** Quantification of VRAC activity in primary astrocyte cultures, measured as swelling-activated D-[^3^H]aspartate release. Data are the mean values ±SEM. n=9 experiments/group performed in three independent cell cultures. ***p<0.001, 4-OHT vs DMSO, The effect of time, treatment, and interaction, repeated measures ANOVA.

Finally, to confirm functional loss of VRAC activity, we employed a radiotracer assay, measuring swelling-activated release of the non-metabolizable glutamate analog, D-[^3^H]aspartate. This assay, which has been thoroughly validated using RNAi and pharmacological tools, quantifies the activity of the LRRC8A-containing VRAC in large populations of astrocytes and gives comparable results to electrophysiological recordings of VRAC Cl^-^ currents (Abdullaev et al., 2006;Hyzinski-Garcia et al., 2014;Wilson et al., 2021). Hypoosmotic challenge of cultured astrocytes stimulated glutamate (d-[^3^H]aspartate) release via VRAC, and such release was reduced by approximately 70% in the tamoxifen-treated iaLRRC8A iKO cultures as compared to the DMSO-treated controls from the same genotype (Figure 2D). Taken together, the results of our *in vivo* and *in vitro* experiments suggest that we successfully downregulated LRRC8A expression in brain astrocytes and that such downregulation was associated with a significant loss of VRAC activity.

### Astrocytic LRRC8A deletion does not confer neuroprotection in an MCAo stroke model

We started with four genotype/treatment groups containing mice of both sexes. Either *Lrrc8a*^fl/fl^ controls or iaLRRC8A KO animals were treated with oil or tamoxifen (see study design in Figure 1A). Ten to 12 days after completion of tamoxifen induction, mice were subjected to 40-min MCAo to investigate potential beneficial effects of eliminating LRRC8A and VRAC in astrocytes. Mice were allowed to survive for 72 h during which time they were evaluated for neurological deficits, followed by quantification of stroke infarction volumes using TTC staining. Because stroke outcomes are significantly impacted by intra- and postischemic variations in blood flow (Hossmann, 1994), we extensively analyzed cerebral blood flow dynamics in real time during the whole MCAo procedure and throughout the first 10 min of reperfusion using a Laser doppler flowmetry (Figure 3A). The laser Doppler probe was placed in the infarction core territory. Twoway ANOVA analysis revealed no significant difference among groups immediately after MCA occlusion as well as near the end of ischemic episode (Figure 3B-C), suggesting similar ischemia severity in all groups. Furthermore, we found no variations in reperfusion rates among all four tested groups (Fig. 3D).

**Figure 3:**
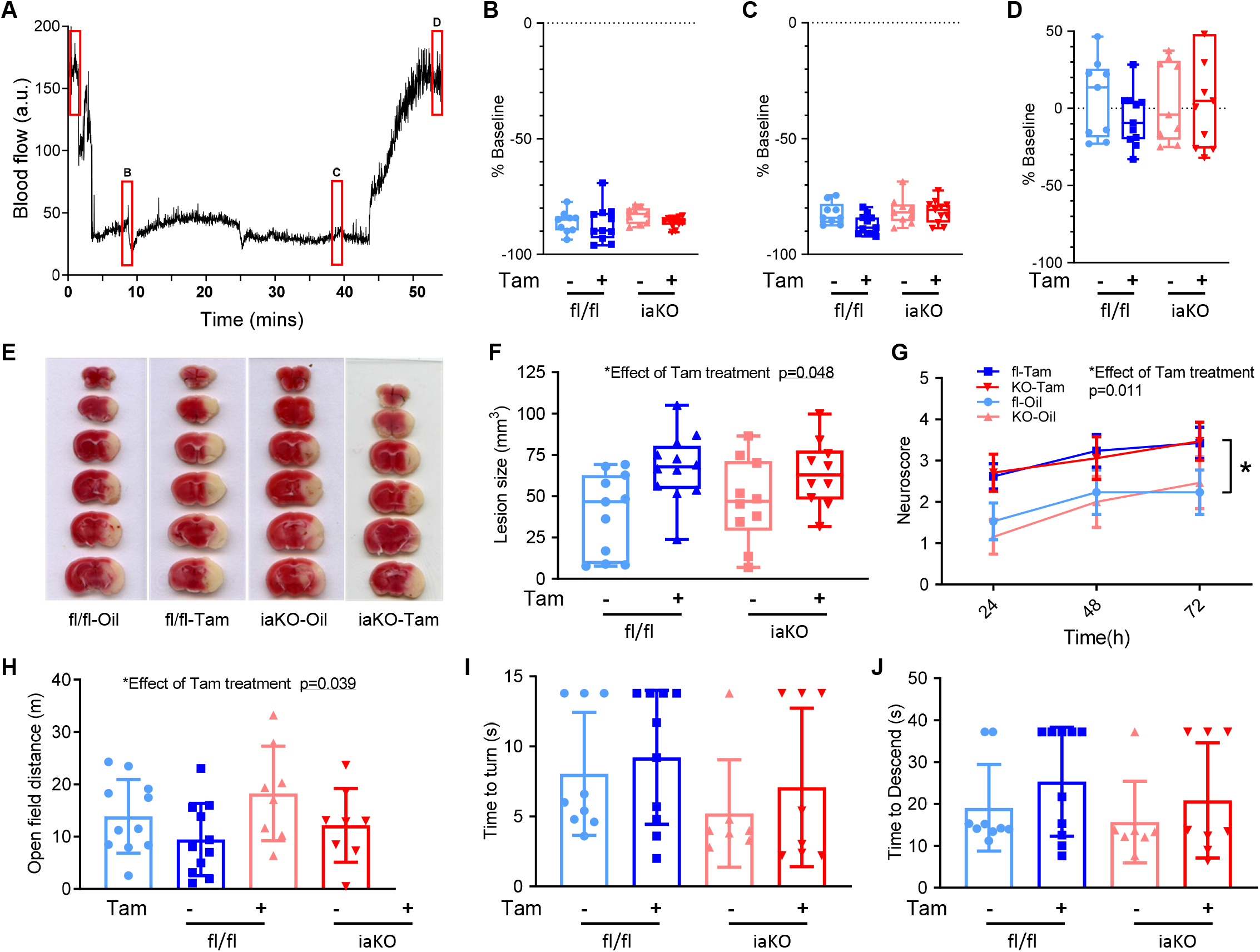
Effects of inducible astrocytic LRRC8A knockout (iaLRRC8A KO) on physiological parameters and outcomes in murine experimental stroke. **(A)** Representative chart of the cerebral blood flow dynamics in experimental stroke as measured with a laser Doppler flowmetry. Cerebral ischemia was induced via occlusion of the middle cerebral artery (MCAo). Red boxes indicate the two-minute intervals used for quantitative analyses of blood perfusion levels. **(B)** The box plot graph of the postocclusion blood flow rates in mice from four genotype/treatment groups (indicated by ‘B’ in the representative graph). n=9-11/group. **(C)** The box plot graph of the pre-reperfusion blood flow levels (‘C’). n=9-11/group. **(D)** The box plot graph of the blood flow recovery during initial reperfusion (‘D’). n=9-11/group. **(E)** Representative TTC staining images of MCAo lesion volumes in four genotype/ treatment groups at 72-h post-stroke. **(F)** Quantitative analysis of MCAo infarction volumes in four experimental groups. *Only the effect of tamoxifen treatment has been found; two-way ANOVA. **(G)** Neurological deficits in four experimental groups assessed daily over 72 h using the neuroscore test. *Only the effect of tamoxifen treatment has been found; repeated measures ANOVA. **(H)** Comparison of spontaneous locomotor activity among groups as assessed by total distance travelled in the open field test. Mean values ±SD. N=8-11. *p<0.05, effect of treatment; two-way ANOVA. **(I)** Lack of effect of treatment and genotype on time to perform a full turn in the pole test. Mean values ±SD. N=8-11. **(J)** Lack of effect of treatment and genotype on time to descend in the pole test. Mean values ±SD. N=8-11.

Our pre-planned primary endpoint was brain lesion size analyzed at 72 h postischemia. In this study, 40-min MCAo produced large infarction in striatal and cortical regions, in line with the previous findings in this model [Figure 3E and (Belayev et al., 1999;Balkaya et al., 2021)]. Two-way ANOVA of brain lesion sizes did not reveal any effect of genotype or interaction between genotype and treatment (Figure 3F). Preplanned post-hoc analysis did not reveal differences between sexes (data not shown). These findings strongly suggest that this strategy of astrocytic LRRC8A deletion is not neuroprotective. Unexpectedly, there was a statistically significant worsening of stroke outcomes after tamoxifen treatment in both genotypes (p=0.005, Figure 3F). The latter finding likely indicates that animals did not fully recover from acute tamoxifen toxicity, even 10-12-day after completion of tamoxifen injections. Overall, we saw no evidence for protection by the astrocyte-specific LRRC8A deletion.

Pre-planned secondary outcomes included post-MCAo mortality and several behavioral assessments of neurological deficits. We found no effect of genotype or genotype-treatment interaction on death rates (data not shown). Our additional secondary endpoint was a neurological deficit score on a five-point scale, which was performed 3 consecutive days after MCAo. We found no effect of genotype or genotype-treatment interaction (Figure 3G). However, on its own, tamoxifen treatment worsened the stroke-induced deficits in a genotype-independent manner (p=0.048, Figure 3G). This was consistent with larger infarctions and reflected the trend for higher mortality in two tamoxifen-treated groups.

We next quantified post-stroke deficits using two sensitive behavioral techniques, the open field test and the pole test, which are commonly employed by the field. The open field test assesses changes in spontaneous activity and anxiety (Balkaya et al., 2013). The pole test is a sensitive measure of simple motor functions, grip strength, and coordination (Balkaya et al., 2013). In the open field test, two-way ANOVA did not reveal the effect of genotype or interaction between treatment and genotype (Figure 3H), corresponding to an absence of beneficial effect of astrocytic LRRC8A deletion. However, yet again, we found the negative effect (reduction in spontaneous locomotion) in two tamoxifen groups, independently of genotype (p=0.039, Figure 3H). In the pole test, two way ANOVA found no effect of genotype, treatment or treatment-genotype interaction on *time to turn* (Figure 3I) and *time to descend* parameters (Figure 3G). Overall, behavioral tests were consistent with histological outcomes and showed no protection by LRRC8A deletion in astrocytes.

### *Nestin*^Cre^-driven deletion of *Lrrc8a* eliminates LRRC8A protein in the brain

The failure to detect histological or behavioral benefits of the astrocyte-specific LRRC8A deletion in stroke, prompted us to re-evaluate the previously reported neuroprotective effects of the brain-wide conditional LRRC8A knockout using *Nestin*^Cre^-driven gene excision. The latter genetic strategy was reported to reduce stroke infarction volumes in LRRC8A-null mice by at least 50% (Zhou et al., 2020).

The *Nestin*^Cre^ knockout strategy allows for effective excision of floxed product(s) in all types of brain neuroectodermal cells, including astrocytes, oligodendrocytes and neurons (Dubois et al., 2006). Nestin is a type VI intermediate filament protein that is abundant in neuronal stem cells throughout development and adulthood; it is expressed in developing mouse embryos as early as E7.5 (Lagace et al., 2007). Complete elimination of LRRC8A in the brain using the *Nestin*^Cre^ strategy has been extensively validated in our recent work (Wilson et al., 2021). In the present study, we used the same breeding strategy (Figure 4A) and additionally used Ai14 tdTomato reporter to ensure that there is no non-specific germline recombination. Our breeding produced four genotypes: control *Lrrc8a*^fl/+^ (containing one wild-type *Lrrc8a* allele; abbreviated fl/+), control *Lrrc8a*^fl/fl^ (fl/fl), heterozygous deletion (*Nestin*^Cre^;*Lrrc8a*^fl/+^, Het), and the full LRRC8A brain knockout (*Nestin*^Cre^;*Lrrc8a*^fl/fl^, hereafter referred to as bLRRC8A KO). We used both sexes of all four genotypes for evaluating the role of LRRC8A protein and VRAC in stroke outcomes.

**Figure 4:**
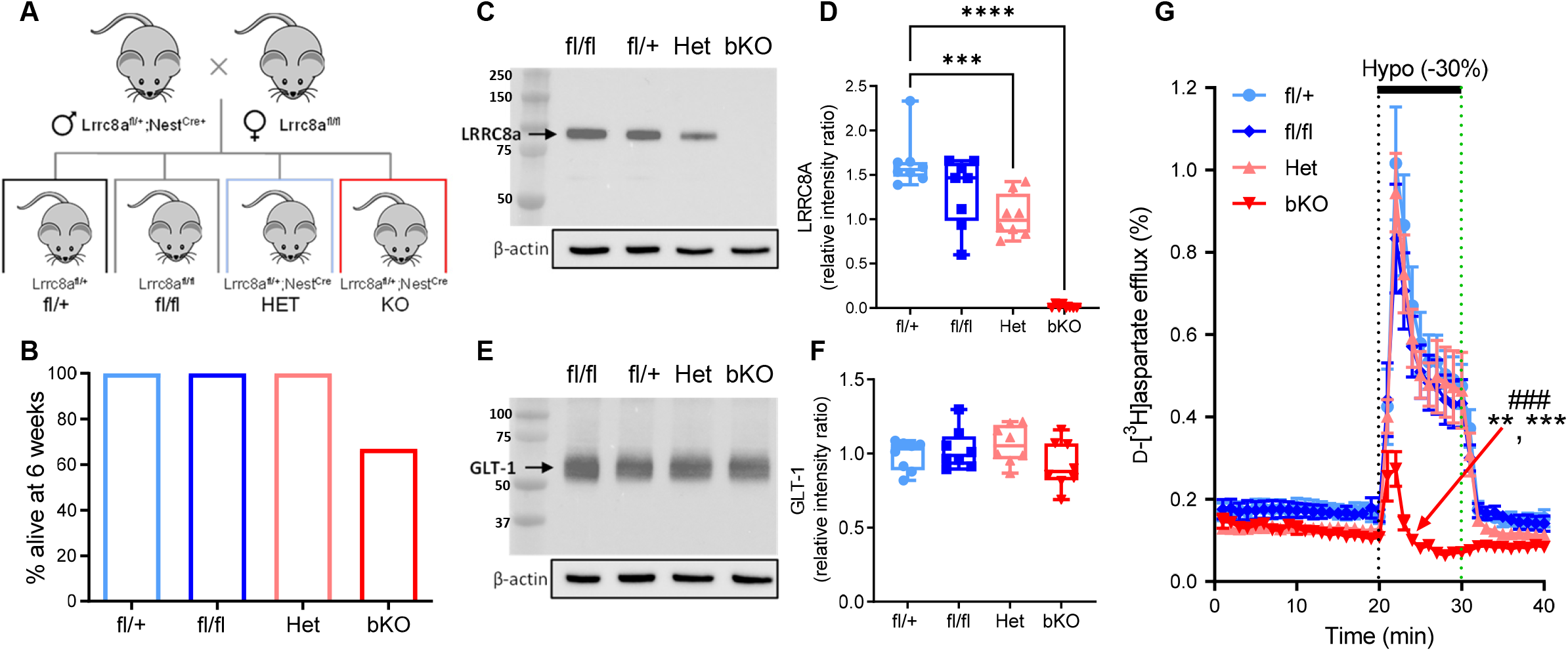
Validation of *Nestin^Cre^* driven LRRC8A deletion *in vivo*. **(A)** Breeding strategy for the conditional deletion of LRRC8A in neurons, astrocytes, and oligodendrocytes using Nestin^Cre^-strategy (bLRRC8A KO). The four potential genotypes from this breeding strategy are *Lrrc8a*^fl/+^ (fl/+), *Lrrc8a*^fl/fl^ (fl/fl), *Nestin*^Cre/+^;*Lrrc8a*^fl/+^ (Het), and *Nestin*^Cre/+^;*Lrrc8a*^fl/fl^ (bKO) mice. **(B)** Survival rates among the four genotypes, with the bKO mice showing > 30% mortality by the week six. **(C)** Representative western blot image of LRRC8A expression in the whole brain lysates from four genotypes. **(D)** Quantification of LRRC8A expression. The box plot of individual IR values, normalized to β-actin loading and averaged within a set. n=8/group. ***p<0.001, Het vs fl/+, ****p<0.0001 bKO vs fl/+. One way ANOVA with Tukey’s multiple comparison test. **(E)** Representative western blot image of GLT1 expression in the whole brain lysates from four genotypes. **(F)** Quantification of GLT1 expression. The box plot of individual IR values, normalized to β-actin loading and averaged within a set. n=8/group. **(G)** Assessment of swelling induced VRAC activity in primary astrocyte cultures, measured as D-[^3^H]aspartate release in response to hypoosmotic medium superfusion. Data are the mean values ±SEM of nine individual experiments per genotype from three different astrocyte cultures. **p <0.01, maximum release, KO versus fl/fl, ***p<0.001, KO vs fl/+ and Het. ^###^p<0.001, integral (10-min) hypoosmotic release values, KO versus all other groups. Reproduced with permission from Wilson et al., 2021.

Much like in our prior work (Wilson et al., 2021) and the study by Zhou et al. (Zhou et al., 2020), bLRRC8A KO mice developed normally but showed spontaneous mortality beginning week 5 with majority of animals dead by week 9. Our prior study established that the main cause of death during adolescence were spontaneous seizures (Wilson et al., 2021). Due to mortality and animal size limitations, we had to perform MCAo procedure at 6-6.5 weeks of age, by which time 40% of bLRRC8A KO animals were already lost (Figure 4B). To ensure that LRRC8A is deleted in the present cohort of animals, we performed semiquantitative western blotting in brain lysates from all four genotypes. Consistent with our prior work (Wilson et al., 2021), we found complete loss of LRRC8A immunoreactivity in bLRRC8A KO animals (Figure 4C, E). Het brains demonstrated ~40% reduction in LRRC8A levels (Figure 4C, E). Interestingly, in fl/fl specimens LRRC8A expression was also reduced by 20% as compared to fl/+ tissue (p=0.093; Figure 4C, E). Although the latter reduction was only a trend, it matched our prior statistically significant findings (Wilson et al., 2021). This phenomenon likely indicates that introduction of loxP sites changes mRNA stability or translation efficacy.

Additionally, we analyzed the expression levels of astrocytic glutamate transporter GLT1. This is important because decreases in GLT1-dependent glutamate buffering may impact stroke outcomes (Rao et al., 2001;Harvey et al., 2011) and we have previously found that the loss of LRRC8A is associated with a partial loss of GLT-1 (Wilson et al., 2021). In the present cohort of animals, GLT-1 levels were not statistically different between the four tested genotypes (Figure 4D, F). The discrepancy between the current and the prior work may be due to the younger age of animals analyzed in the current study.

### VRAC activity is lost in bLRRC8A KO astrocytes but not affected by heterozygous bLRRC8A deletion

The functional loss of VRAC activity in bLRRC8A KO has been extensively characterized in our prior work (Wilson et al., 2021) (reproduced with permission in Figure 4G). The critical finding of this latter study is nearly complete loss of swelling-activated glutamate (d-[^3^H]aspartate) release in bLRRC8A KO astrocytes (Figure 4G). The small residual d- [^3^H]aspartate efflux present in the KO cultures displays a different time-dependent inactivation kinetics and may be mediated by an alternative-to-VRAC release pathway. Equally important, astrocytic cultures derived from Het brains had virtually identical VRAC-mediated glutamate release as compared to fl/+ and fl/fl astrocyte controls (Figure 4G). This suggests that 20% and 40% reductions in LRRC8A protein levels in fl/fl and Het astrocytes, respectively, are insufficient to reduce plasmalemmal VRAC levels and activity. Taken together, these results confirmed the successful deletion of LRRC8A in bKO mice and the full functional VRAC activity in bLRRC8A Het animals.

### Both complete and heterozygous bLRRC8A deletion reduces stroke damage

We independently reproduced the recent study by Zhou et al. (Zhou et al., 2021) measuring stroke infarction volumes in bLRRC8A KO mice. One important addition in the present work, was the use of four genotypes (fl/+, fl/fl, Het, and KO) rather than a straightforward comparison of WT to bLRRC8A KO animals in the aforementioned publication. As it will be apparent from further discussion, the four-genotype comparison allowed for deeper interpretation of obtained results. Because of mortality in bLRRC8A KO mice, we used 6-6.5-week-old animals, which were subjected to a 40-min MCAo with reperfusion. To make our results directly comparable to the published work, we quantified stroke lesion volumes at 24 h rather than 72 h.

For iaLRRC8A KO experiments, we started with the comparisons of cerebral blood flow dynamics during and after MCAo (representative graphs in Figure 5A). As expected, we found the strong and persistent drop in blood during MCAo (Figure 5A). One way ANOVA found no differences among four genotypes immediately after occlusion and at the end of MCAo episode (Figure 5B, C). In the initial stages of reperfusion, blood flow values returned close to the pre-MCAo levels, with the exception of bLRRC8A KO group showing ~20% blood flow deficit (representative chart and average values in Figure 5A, D). This trend prompted us to partially “relax” our pre-planned inclusion criterion for reperfusion. The post-hoc analysis indicated that the reperfusion deficit in bLRRC8A KO mice was statistically different from fl/fl group only (p=0.012, Figure 5D). Overall, these data suggested that we have similar severity of ischemia in all genotypes, and that stroke outcomes in bLRRC8A KO group could be worsened by deficits in early reperfusion.

**Figure 5:**
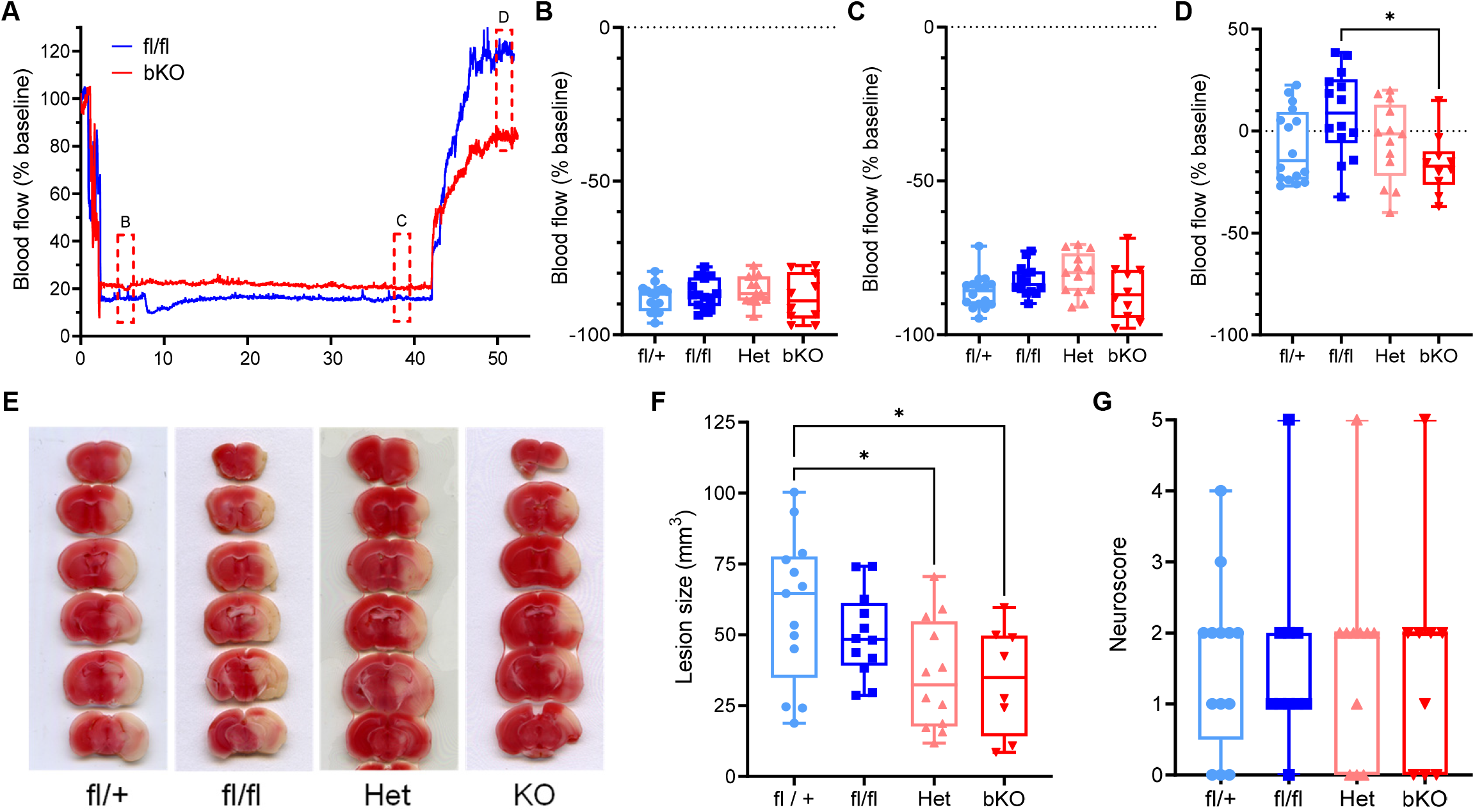
Effects of conditional brain-wide LRRC8A knockout (bLRRC8A KO) on primary and secondary outcomes in murine experimental stroke. **(A)** Representative changes in cerebral blood flow levels in bKO and control fl/fl mice during an MCAO experiment, measured using a laser Doppler flowmetry. **(B)** The box plot graph of the post-occlusion blood flow rates in mice from four genotype groups (indicated by ‘B’ in the representative graph). n=10-16/group. **(C)** The box plot graph of the pre-reperfusion blood flow levels (‘C’). n=10-16/group. **(D)** The box plot graph of the blood flow recovery during initial reperfusion (‘D’). n=10-16/group. **(E)** Representative TTC staining images of MCAo lesion volumes measured at 24 h post stroke. **(F)** Box plot graph and analysis of the primary outcome, the stroke infarction volumes corrected for brain edema, among four genotypes. *p<0.05, bKO vs fl/+, Het vs fl/+. One-way ANOVA with Tukey’s correction for multiple comparisons. n=8-13/ group **(G)** Quantitative comparison of neurological deficits in four genotypes measured at 24 h post-stroke as assessed by the five-point Neuroscore test.

Our pre-planned primary outcome was the ischemic lesion size measured with a TTC approach. The 40-min MCAo produced large striatal and cortical infarction (representative images in Figure 5A), which highly resembled the outcomes in older animals included in iaLRRC8A KO analysis (compare to Figure 3E). One way ANOVA of lesion sizes found that bLRRC8A KO and Het mice had significantly smaller brain infarctions as compared to the fl/+ group, with the reduction of ~50% (Figure 5F). However, when the same two groups were compared against fl/fl controls, the neuroprotection was largely lost (Figure 5F). One-way ANOVA comparisons of the secondary outcome, the neurological deficit on the five-point Neuroscore scale, did not reveal any differences among groups (Figure 5B). Overall, the results of stroke experiments in bLRRC8 KO mice and littermate controls showed a lack of correlation between infarction volumes and swelling-activated glutamate release.

### Marginal effect of the VRAC blocker DCPIB on intraischemic glutamate release

To gain an additional insight into the mechanism of intraischemic glutamate release, we performed a microdialysis study quantifying extracellular levels of glutamate, aspartate, and taurine in stroke penumbra. The relatively large size of microdialysis probe (~2 mm) in relationship to small size of mouse brain does not allow for selective sampling of amino acids in penumbra. This is not a small matter because mechanisms of glutamate release fundamentally differ among ischemic regions. In the ischemic core, glutamate release is dominated by the reversal of glutamate transporters (Phillis et al., 1994;Seki et al., 1999) while in the ischemic penumbra it is thought to be mediated by VRAC channels (Feustel et al., 2004;Zhang et al., 2008). For this reason, we conducted microdialysis experiments in a rat model of stroke, as in our prior work (Dohare et al., 2014).

As a positive control for VRAC-mediated glutamate release, we tested the effect of DCPIB on glutamate levels in the rat cortex perfused with hypoosmotic aCSF. We and others validated this approach for measuring VRAC activity *in vivo* [e.g. (Haskew-Layton et al., 2008)]. As shown in Figure 6, perfusion of hypoosmotic solution triggered transient release of glutamate and two other VRAC-permeable amino acids, aspartate and taurine (Figure 6C-E). When hypoosmotic medium was supplemented with the VRAC blocker DCPIB, stimulation of glutamate, aspartate and taurine release was inhibited by 71%, 74%, and 72% respectively (Figure 6C-E). Altogether these experiments confirmed that cellular swelling dramatically upregulated release of VRAC-permeable amino acids, and that the chosen mode of DCPIB delivery is effective in limiting this process.

**Figure 6:**
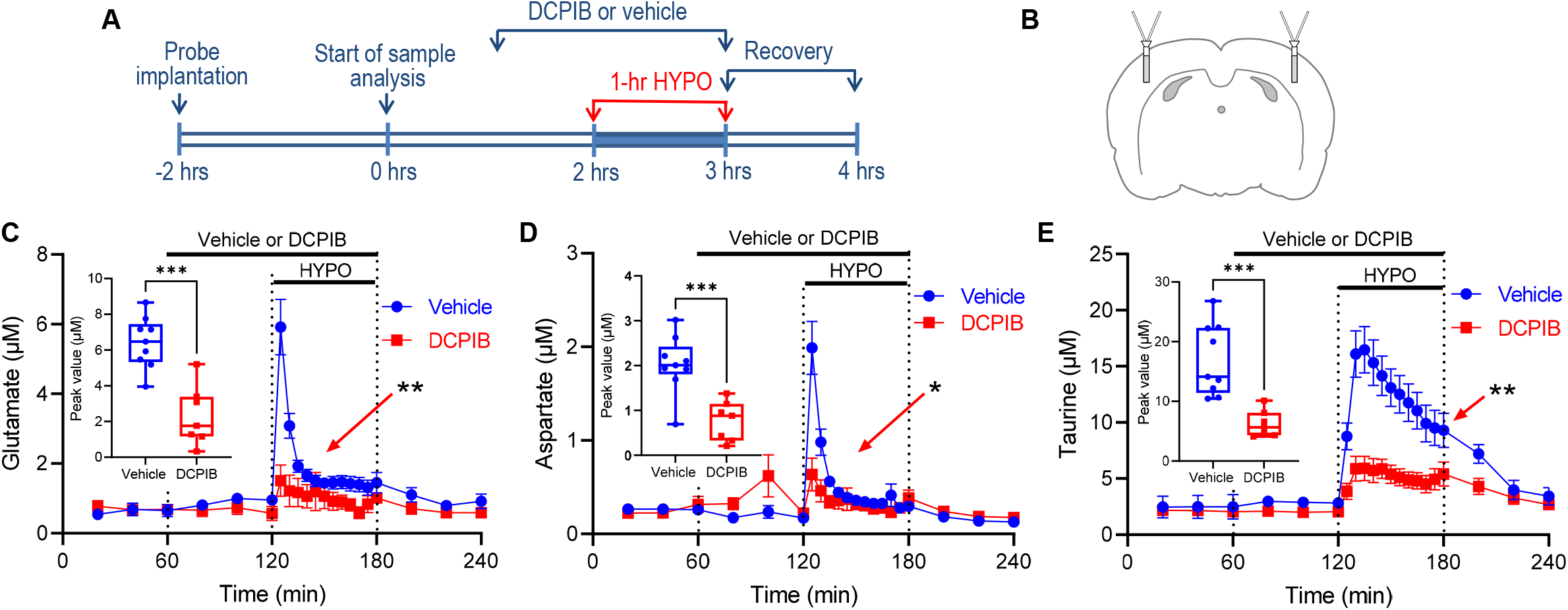
The effects of the VRAC blocker DCPIB on swelling-activated amino acid release *in vivo*. **(A)** Design of microdialysis experiments performed to test the efficacy of the VRAC inhibitor DCPIB *in vivo*. **(B)** Schematic representation of microdialysis probe placements in the rat brain. **(C-E)** The effect of hypoosmotic medium on the extracellular levels of glutamate **(C)**, aspartate **(D)**, and taurine **(E)** in microdialysis samples measured with an HPLC assay. In the C-E fields: *p<0.05, **p<0.01, DCPIB-vs. vehicle-treated groups, the effects of treatment, time, and group-time interaction as analyzed with the mixed-effects model with Geisser-Greenhouse correction. Data are the mean values ±SEM, n=6-9/group. The box-plot insets show the peak values for individual amino acids during the hypoosmotic challenge. ***p<0.001, DCPIB vs Vehicle, t-test.

In the main set of experiments presented in Figure 7, microdialysis probes were placed in the same cortical areas as in hypoosmotic experiments, but now corresponded to ischemic penumbra or its anatomical equivalent in the contralateral hemisphere. The proper positioning of microdialysis probe in the penumbra was confirmed by a laser Doppler flowmetry as shown in Figure 7B. In both treatment groups, the initial reduction of cerebral perfusion was identical; however, the DCPIB-treated animals showed a trend for partial blood flow recovery during the middle cerebral artery occlusion period (p=0.094 for group effect; p=0.208 for group-time interaction; repeated measures ANOVA). Although this effect was not statistically significant, it might be relevant to changes in amino acid levels (see below). The two-hour occlusion of the MCA led to a dramatic increase in the extracellular levels of VRAC permeable glutamate (Figure 7D), aspartate (Figure 7E), and taurine (Figure 7F). Perfusion with DCPIB marginally reduced the intraischemic release of glutamate, aspartate, and taurine by 24%, 24%, and 26%, respectively (Figure 7D-F, insets). Such a reduction reached the level of statistical significance for taurine only. Overall, these results suggest small-to-negligible contribution of VRAC to the intraischemic amino acid release. Also, we cannot exclude that the effect of DCPIB was due to small improvements in the intraischemic blood flow (see Figure 7C) rather than direct inhibition of VRAC.

**Figure 7:**
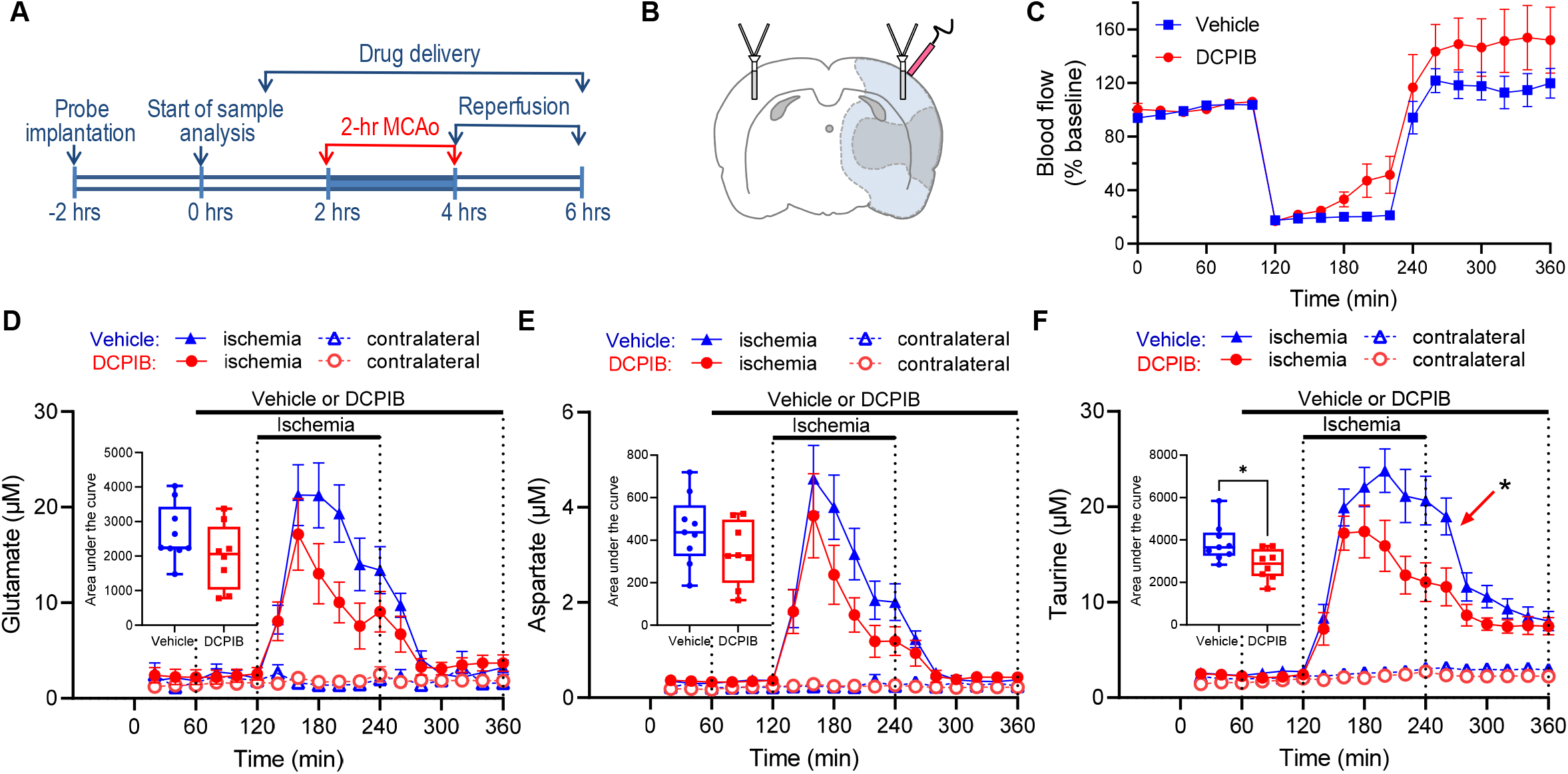
Marginal effects of the VRAC blocker DCPIB on the intraischemic amino acid release *in vivo*. **(A)** Design of microdialysis experiments performed to test the effects of the VRAC inhibitor DCPIB on pathological amino acid release in the rat MCAo stroke. **(B)** Schematic representation of microdialysis and laser Doppler probe placements in the ischemic penumbra and corresponding contralateral location. **(C)** The comparison of blood flow dynamics in Vehicle-and DCPIB-treated groups during and after MCAo, measured using a laser Doppler technique located as indicated in (B). **(D-F)** The effect of DCPIB on the pathological extracellular levels of glutamate **(D)**, aspartate **(E)**, and taurine **(F)** in microdialysis samples collected during MCAo. In the F field: *p<0.05, DCPIB-vs. vehicle-treated taurine levels, the effects of treatment, time, and group-time interaction as analyzed with the mixed-effects model and Geisser-Greenhouse correction. Data are the mean values ±SEM, n=8-9/group. The box-plot insets show the integral release values for individual amino acids collected during the entire MCAo episode. *p<0.05, DCPIB vs Vehicle, for taurine only, t-test.

## DISCUSSION

The major finding of our work is the disconnect between stroke outcomes and relative changes in VRAC activity in two mouse lines with targeted deletion of LRRC8A in brain cells (either in astrocytes only [iaLRRC8A KO] or neurons, astrocytes, and oligodendrocytes [the “brain-wide” bLRRC8A KO]). This surprising discovery is at odds with the widely accepted idea that multifunctional VRAC channels propagate excitotoxic tissue damage via pathological release of the excitatory neurotransmitters, glutamate and aspartate. The main argument against VRAC contributions to glutamate toxicity is the similar reduction of stroke infarction in bLRRC8A KO mice carrying full LRRC8A deletion and their heterozygous littermates. Despite equal protection, bLRRC8A KO rodents have no functional VRAC in brain cells, while heterozygous mice exhibit full VRAC activity. These results strongly suggest that LRRC8A protein may promote ischemic brain damage via a mechanism(s) that is (are) unrelated to glutamate release.

Our first series of experiments tested the protective effect of inducible LRRC8A deletion in astrocytes (iaLRRC8A KO) using *Aldh1l1*^CreERT2^. Astrocytes show the most prominent swelling in stroke and are thought to be the main source of pathological VRAC activity in the CNS (Mongin and Kimelberg, 2005;Kimelberg, 2005;Wilson and Mongin, 2018). Despite strong reductions in astroglial LRRC8A and VRAC-mediated glutamate release, we failed to detect predicted histological or behavioral protection in iaLRRC8A KO, as compared to three genotype/treatment controls. The lack of protection in our study differed from a ~30% reduction in the ischemic brain damage identified in a recent publication which produced a non-inducible LRRC8A deletion in astrocytes (Yang et al., 2019). In the latter work, the authors used the *GFAP*^Cre^-driven approach, which targets neural precursors and astrocytes through all stages of development (Yang et al., 2019). This may be associated with the risk of developmental compensation. Previously, we found that the *Nestin*^Cre^-driven, brain-wide LRRC8A KO causes substantial biochemical alterations in the brain and results in 100% animal mortality in early-to-late adolescence (Wilson et al., 2021). Importantly, in this latter model, we also discovered robust changes in the LRRC8A-null astrocytes, including reactive astrogliosis and downregulation of several transporters and enzymes involved in neurotransmitter homeostasis (Wilson et al., 2021). Such changes appear to be avoided in the current iaLRRC8A KO mice. For example, we do not see reactive astrogliosis or downregulation of astrocytic glutamate transporter GLT-1, both of which were reported in the *Nestin*^Cre^-driven LRRC8A KO mice. These caveats notwithstanding, the combined results of our work and those in the study by Yang et *al*. (Yang et al., 2019) suggest that astrocytic LRRC8A and VRAC play a minor role in ischemic stroke pathology.

The lack of neuroprotection in the iaLRRC8A KO contradicts the prevalent idea that astrocytic VRAC is the key player in stroke pathology. This notion was introduced in the 1990s, based on the findings that hypoosmotic or high K^+^-induced swelling of astrocytes opens an anion permeability pathway and triggers release of the excitotoxic glutamate and aspartate (Kimelberg et al., 1990;Rutledge and Kimelberg, 1996).

Because astroglia are known to be prominently swollen in stroke, it has been suggested that cell volume-sensitive glutamate release contributes to ischemic brain damage (Kimelberg and Mongin, 1998;Kimelberg, 2005;Mongin, 2007). Accordingly, in rodent models of stroke, systemic delivery of the non-selective VRAC blocker tamoxifen decreased brain infarction volumes by as much as 60-80% (Kimelberg et al., 2000;Kimelberg et al., 2003;Mehta et al., 2003;Feng et al., 2004). The tamoxifen protection was seen in adult and neonatal rats, confirmed in both transient and permanent cerebral ischemia, and further corroborated in large animals, in the thromboembolic stroke model in dogs (Kimelberg et al., 2000;Kimelberg et al., 2003;Mehta et al., 2003;Feng et al., 2004;Boulos et al., 2011). Together with subsequent reports of protection with a more selective VRAC inhibitor DCPIB (Zhang et al., 2008;Alibrahim et al., 2013), these studies cemented the notion of VRAC as a promising target for stroke intervention.

Despite seemingly robust evidence for the pathological role of VRAC, the major limitation of prior pharmacological studies is the numerous off-target effects of both tamoxifen and DCPIB. In the therapeutically relevant range of 1–10 μM, tamoxifen blocks diverse voltage-gated and ligand gated Na^+^ and cation channels (Allen et al., 1998;Hardy et al., 1998) or, alternatively, activates the BK subtype of K^+^ channels (He et al., 2003). It also inhibits neuronal and inducible isoforms of nitric oxide synthase and possesses significant antioxidant properties (Osuka et al., 2001;Custodio et al., 1994). On their own, these off-target effects could be neuroprotective, complicating interpretation of stroke findings. Likewise, the more specific DCPIB blocks several pathologically relevant glutamate release pathways and directly activates the two-pore domain and BK K^+^ channels (Bowens et al., 2013;Minieri et al., 2013;Zuccolini et al., 2022). Thus, the exact mechanism for DCPIB protection is also not certain.

The caveats of pharmacological findings have been partially addressed by analyzing stroke outcomes in mice carrying conditional knockout of the essential VRAC subunit LRRC8A. The limited protection conveyed by astrocytic LRRC8A deletion seen in the prior publication (Yang et al., 2019) and the lack of protection in the current work are at odds with the potent reduction of ischemic damage by tamoxifen and DCPIB. Interestingly, another publication found more robust (~50%) reduction of stroke damage in mice carrying the brain-wide LRRC8A knockout produced with *Nestin*^Cre^ (Zhou et al., 2020). Perhaps, these latter results suggest that under pathological conditions, VRAC is active in numerous brain cell types. To definitively examine this hypothesis, we reevaluated stroke outcomes in the *Nestin*^Cre^-driven, brain-wide LRRC8A KO mice. These mice show the complete loss of brain LRRC8A immunoreactivity, electrophysiological VRAC currents, and swelling-activated glutamate release (Yang et al., 2019;Zhou et al., 2020;Wilson et al., 2021). Using this genetic strategy, Zhou and coworkers compared cerebral infarction in the two genotypes, *Lrrc8a*^fl/fl^ controls and LRRC8A KO (Zhou et al., 2020). For additional rigor, we tested stroke outcomes in four distinct genotypes (two controls, *Lrrc8a*^fl/+^ and *Lrrc8a*^fl/fl^, heterozygous *Lrrc8a* deletion, and full *Lrrc8a* KO). Such extended analysis proved to be critical. Consistent with the prior publication, we found that the LRRC8A KO brains were strongly protected against stroke as compared to fl/+ littermates, with the reduction in infarction volumes reaching 50%. Yet, when LRRC8A KO results were analyzed against fl/fl controls, the average protection was smaller and not statistically significant. However, the key finding came from the assessment of LRRC8A Het mice. *LRRC8A KO and Het animals were equally protected against stroke, even though LRRC8A Het cells retained full VRAC activity*. In our judgement, a complete disconnect between VRAC activity and stroke outcomes rules out significant role for the VRAC-mediated glutamate release in acute ischemic stroke injury.

To additionally probe the pathological contributions of VRAC we employed a microdialysis approach and measured glutamate levels in the ischemic penumbra. Because of the smaller size of the mouse brain, these experiments were conducted in rats. As with several prior studies, we found dramatic elevation in the extracellular levels of glutamate and the other VRAC permeable neurotransmitters, aspartate and taurine. Yet, in our hands, the VRAC blocker DCPIB showed minor impact on pathological amino acid release. Together with the lack of correlation between swelling-activated glutamate release and neuroprotection in our genetically modified animals, these results support the conclusion that VRAC-mediated glutamate release does not dominate excitotoxic tissue damage in the ischemic penumbra. The apparent lack of VRAC activity in stroke tissue may be explained by the requirement of non-hydrolytic ATP binding for VRAC channel opening (Strange et al., 1996;Okada, 1997;Nilius et al., 1997). Metabolic inhibition or omission of ATP from pipette solution eliminate swelling-activated Cl^-^ currents and amino acid release (Jackson et al., 1994;Oike et al., 1994;Oiki et al., 1994;Rutledge et al., 1999). This high dependence on cytosolic ATP or other nucleotide triphosphates is likely to limit VRAC activity in metabolically compromised ischemic tissue, particularly in non-astroglial cells and explain our microdialysis results.

If swelling-activated glutamate release is not involved, then why are LRRC8A Het and KO brains protected against stroke? One speculation is that changes in LRRC8A protein expression may modify pro-death intracellular machinery in neural cells. Besides its essential role in the VRAC channel function (Qiu et al., 2014;Voss et al., 2014), LRRC8A protein additionally serves as a signaling scaffold molecule. Thus, LRRC8A physically interacts with two important adaptor molecules, Growth factor Receptor-Bound protein 2 (GRB2) and GRB2-Associated Binding protein 2 (GAB2), and in such a way potently modulates downstream signaling cascades and modifies many cellular functions (Kumar et al., 2014;Zhang et al., 2017;Kumar et al., 2020;Alghanem et al., 2021). Among the relevant pathways, LRRC8A expression seems to promote activities of c-Jun N-terminal kinases (JNK) (Choi et al., 2016;Lu et al., 2019). JNK isoforms, and particularly JNK3 promote ischemic/hypoxic brain injury and their acute inhibition protects against stroke damage in mice (Kuan et al., 2003;Borsello et al., 2003;Gao et al., 2005;Zulfiqar et al., 2020). It is conceivable that LRRC8A-dependnet changes in this or other parts of intracellular machinery could reduce stroke damage

The practical implication of our work is that the utility of *selective* inhibition of VRAC for reducing ischemic brain damage needs to be reevaluated. Over the years, the field accumulated strong evidence for neuroprotective actions of the putative VRAC blockers, tamoxifen and DCPIB. Yet, the protection by these agents appears to be multifactorial and largely unrelated to the VRAC-mediated glutamate release. Likewise, stroke protection in mice with targeted deletion of the essential VRAC subunit LRRC8A, may also be unrelated to glutamate toxicity and engage distinct molecular processes.

## MATERIALS AND METHODS

### Key resources table

Key resources table:
| Reagent type (species) or resource | Designation | Source or reference | Identifiers | Additional Information |
| --- | --- | --- | --- | --- |
| Strain, strain background (M. musculus) | *Lrrc8a*<sup>fl/fl</sup> (*Swell1*<sup>fl/fl</sup>) mice (C57BL/6J background) | PMID: 28436964 |  | Shared by Dr. Rajan Sah |
| Strain, strain background (M. musculus) | B6N.FVB-Tg (Aldh1l1-cre/ERT2)1Khakh/J | Jackson Laboratory | Stock # 031008 |  |
| Strain, strain background (M. musculus) | *Aldh*<sup>CreERT2/+</sup>; *Lrrc8a*<sup>fl/fl</sup> | This paper |  | Astrocyte-specific inducible LRRC8A knockout (aiKO) |
| Strain, strain background (M. musculus) | (B6.Cg-Tg(Nes-cre)1<sup>Kln</sup>/J | Jackson Laboratory | Stock #003771 |  |
| Strain, strain background (M. musculus) | *Nestin*<sup>Cre/+</sup>; *Lrrc8a*<sup>fl/+</sup> | This paper |  | Heterozygous deletion of LRRC8A (HET) |
| Strain, strain background (M. musculus) | *Nestin*<sup>Cre/+</sup>; *Lrrc8a*<sup>fl/fl</sup> | This paper |  | Brain-specific LRRC8A knockout (bKO) |
| Cell line (M. musculus) | Cultured primary astrocytes | This paper |  | Prepared from six genotypes of iaKO and bKO mice |
| Antibody | Mouse monoclonal glial fibrillary acidic protein (GFAP) | Millipore-Sigma | cat. #MAB360 | IHC dilution: 1:400 |
| Antibody | Mouse monoclonal anti-glutamine synthetase (GS) | Millipore-Sigma | cat. # G2781 | IHC dilution: 1:5,000; RRID:AB_259853 |
| Antibody | Mouse monoclonal anti-GLT1(EAAT2) | Abcam | cat. # AB1783 | WB dilution: 1:25,000 |
| Antibody | Mouse monoclonal anti-LRRC8A | Santa Cruz | cat. # sc-517113 | WB dilution: 1:500 |
| Antibody | Donkey polyclonal anti-rabbit IgG, horseradish peroxidase (HRP)-conjugated antibody | GE Healthcare | cat. #NA932 | Secondary Ab, WB dilution: 1:10,000 |
| Antibody | HRP-conjugated sheep anti-mouse IgG | GE Healthcare | cat #NA931 | Secondary Ab, WB dilution: 1:10,000 |
| Antibody | HRP-conjugated primary anti-β-actin antibody | Millipore-Sigma | cat. #A-3854 | WB dilution 1: 50,000 |
| Antibody | goat anti-mouse Alexa-Fluor 488 | ThermoFisher Scientific | cat # A32723 | IHC dilution: 1:400 |
| Antibody | goat anti-rabbit Alexa-Fluor 647 | ThermoFisher Scientific | cat. #A21244 | IHC dilution: 1:400; RRID:AB_2535812 |
| Antibody | donkey anti-mouse Alexa-Fluor 594 | ThermoFisher Scientific | cat #A-32744 | IHC dilution: 1:400 |
| Antibody | goat anti-rabbit Alexa-Fluor 488 | ThermoFisher Scientific | cat # A32731 | IHC dilution: 1:500 |
| Chemical reagent | Corn oil | Millipore-Sigma | cat # C8267 | Vehicle for i.p. injections |
| Chemical reagent | 4-Hydroxy-tamoxifen | Millipore-Sigma | cat # H6278 | Cre induction *in vitro* |
| Chemical reagent | Tamoxifen | Millipore-Sigma | cat # T5648 | Cre induction *in vivo*, 120 mg/kg |
| Chemical reagent | ProLong Glass Antifade Mountant | ThermoFisher Scientific | cat. # P36984 |  |
| Chemical reagent | Protease inhibitor cocktail | ThermoFisher Scientific | cat. # 78430 |  |
| Chemical reagent | Laemmli buffer | Bio-Rad | cat. # 1610737 |  |
| Chemical reagent | Paraformaldehyde, 4% in PBS | Santa Cruz Biotechnology | cat. # sc-281692 |  |
| Chemical reagent | Restore buffer | ThermoFisher Scientific | cat. #21059 |  |
| Chemical reagent | Tissue-Tek O.C.T. compound | Sakura Finetek | cat.#4583 |  |
| Commercial assay or kit | ECL reagent | GE Healthcare | cat. #RPN2232 |  |
| Commercial assay or kit | Bicinchoninic acid assay | ThermoFisher Scientific | Cat.# 23225 |  |
| Software | NeuroLucida 11.03 | MBF Bioscience |  | RRID:SCR_016788 |
| Software | ANY-maze | Stoelting Co. |  | Video tracking software<br>RRID:SCR_014289 |
| Software | Prism 7.0 | GraphPad, San Diego, CA, USA |  | RRID:SCR_002798 |

### Ethics statement

All animal procedures used in the present study were approved by the Institutional Animal Care and Use Committee of Albany Medical College (ACUP 18-04002, 18-12001, 18-12003, 21-11002), and strictly conformed to the Guide for the Care and Use of Laboratory Animals as adopted by the U.S. National Institutes of Health (https://grants-nih-gov.elibrary.amc.edu/grants/olaw/Guide-for-the-Care-and-Use-of-Laboratory-Animals.pdf).

### Animals

Mice were housed in a temperature, humidity, and light-controlled facility on a 12:12 h light/dark cycle and given free access to food and water. Experiments were performed in age-matched controls of both sexes, divided as evenly as possible between males and females. *Lrrc8a*^fl/fl^ (*Swell1*^fl/fl^) mice were generated as previously described (Zhang et al., 2017).

Astrocyte-specific, inducible *Lrrc8a* knockout mice were produced by breeding *Lrrc8a*^fl/fl^ female mice with commercially available *Aldh*^CreERT2/+^ male mice (B6N.FVB-Tg(Aldh1l1-cre/ERT2)1Khakh/J, stock # 031008) both on a C57BL/6J background. After initial steps, *Aldh1l1*^Cre/+^;*Lrrc8a*^fl/fl^ heterozygous males were crossed again with *Lrrc8a*^fl/fl^ females mice to produce *Aldh1l1*^Cre/+^;*Lrrc8a*^fl/fl^ inducible astrocytic knockout mice and *Lrrc8a*^fl/fl^ littermate controls. To activate Cre ice were treated with either tamoxifen (120 mg/kg, five daily injections) or an equal volume of vehicle (corn oil).

Brain-specific *Lrrc8a* knockout mice were produced by breeding *Lrrc8a*^fl/fl^ female mice with commercially available *Nestin*^Cre/+^ male mice (B6.Cg-*Tg(Nes-cre)1^Kln^*/J; Jackson Laboratory stock #003771), both on a C57BL/6 background. *Nestin*^Cre/+^;*Lrrc8a*^fl/+^ heterozygous males were crossed again with *Lrrc8a*^fl/fl^ female mice to produce *Nestin*^Cre/+^;*Lrrc8a*^fl/fl^ knockout mice and littermate controls.

All genotypes were confirmed by PCR analysis across the predicted loxP insertion sites surrounding *Lrrc8a e*xon 3, or for the *Aldh1l1*^Cre^ and *Nestin*^Cre/+^ transgene insertion according to the Jackson Laboratory genotyping protocols.

All microdialysis studies were performed in male Sprague-Dawley rats (Taconic Farms, 230-280 g). Animals were maintained on a 12/12-h light/dark cycle and allowed free access to food and water.

### Cell cultures of brain astrocytes

Primary cultures of mouse astrocytes were prepared from cortices of newborn (P0-P1) male and female mice. Complete litters were utilized, and tail snips of each pup were taken for genotyping. Neonates were anesthetized by cooling and rapidly decapitated. Brains were harvested in ice-cold sterile Dulbecco’s phosphate-buffered saline without calcium and magnesium (ThermoFisher Scientific, cat. #14190144). Cortical tissue was dissected from meninges, minced using a sterile surgical blade, and additionally triturated by pipetting. Triturated tissue was then digested for 5 min using the recombinant protease TrypLE Express (ThermoFisher Scientific, cat. #12605) at room temperature. The dissociated cells were sedimented at 900 g for 10 minutes at 4°C. Cell pellets were resuspended in Earl’s minimal essential medium (MEM) supplemented with 10% heat-inactivated horse serum (HIHS) and penicillin/streptomycin (all components from ThermoFisher Scientific/Invitrogen). To obtain an enriched astrocyte population, cells from each individual brain were separately plated onto an T75 flask pretreated with poly-D-lysine (Sigma-Millipore, cat. #P6407) and grown in a humidified atmosphere of 5% CO2/balance air at 37°C. Cell cultures typically reached confluency around 10 days and were maintained up to six weeks until replating for functional assays. The purity of astrocyte populations was periodically verified with immunocytochemistry by staining for the astrocyte marker, glial fibrillary acidic protein (GFAP, Millipore-Sigma, 1:400), and exceeded 90%.

### Processing of brain tissue samples

To collect brain tissue, mice were transcardially perfused with either PBS for western blot, or 4% paraformaldehyde in PBS for immunohistochemistry after complete anesthesia with a lethal dose of sodium pentobarbital (100 mg/kg).

For immunohistochemistry, brains were removed and additionally postfixed in 4% paraformaldehyde for 24 h, dehydrated in 15% sucrose solution for 24 h, and then in 30% sucrose solution for full cryopreservation (all steps at 4°C). Brains were attached to a cryostat specimen stage using Tissue-Tek OCT compound, allowed to freeze to −20°C and sectioned using a CM3050 cryostat (Leica, Wetzlar, Germany, RRID: SCR_016844) into 25-μm thick sections. Sections were placed into cryoprotectant media containing 30% ethylene glycol and 20% glycerol in Tris-buffered saline (TBS) and stored at −20°C until immunohistochemistry is performed.

For western blotting, brains were removed, sectioned to two hemispheres and immediately snap-frozen in liquid nitrogen. A single hemisphere was homogenized using a mechanical homogenizer (PRO200, PRO Scientific, Oxford, CT, USA) in a tissue lysis buffer containing 150 mM NaCl, 50 mM Tris HCl (pH 8.0), 0.1% Triton X-100, and 5% protease inhibitor cocktail. Samples were clarified by a brief centrifugation (10,000 *g* for 5 min at 4°C) and the supernatants were collected and diluted with 2× reducing Laemmli buffer.

### Immunohistochemistry

For immunohistochemical detection, floating sections were washed with PBS and blocked in PBS with 2% normal goat serum and 0.2% TX-100 for one hour at room temperature. They were next incubated overnight at 4°C in the same solution additionally containing mouse monoclonal anti-GFAP antibody (1:400) and rabbit polyclonal anti-glutamine synthetase antibody (1:5,000). The next morning, brain sections were washed in PBS and incubated for 2 h at room temperature with the secondary antibodies: goat antimouse Alexa-Fluor 488 (1:400 for GFAP) and goat anti-rabbit Alexa-Fluor 647 (1:400 for GS). After PBS wash, sections were counterstained with 0.05 μg/ml DAPI, additionally washed with PBS, and transferred onto glass slides. Slides were dried overnight at room temperature in the dark, and then mounted with ProLong Glass Antifade Mountant. Immunolabeled brain sections were scanned using a Zeiss AX10 microscope at 200× magnification, and the resulting images stitched together with Neurolucida 11.03 software. Alternatively, to determine 3D-colocalization of tdTomato reporter and astrocytic makers, images were acquired using a Zeiss LSM880 confocal microscope with a LCI Plan-Neofluar 25×/0.8 Imm Korr DIC objective (Carl Zeiss, RRID:SCR_0113672).

### Western blotting analysis

The success of LRRC8A deletion was confirmed using semi-quantitative Western blot analyses in protein lysates from brain tissue and astrocyte cultures. Additionally, we quantified changes in the brain expression levels of the astrocytic glutamate transporter GLT-1. Protein lysates in the reducing Laemmli buffer were boiled for 5 min, loaded onto 10% Mini-PROTEAN TGX gels (Bio-Rad, cat #4561033), and separated according to a standard SDS-PAGE protocol. Separated proteins were electrotransferred onto a polyvinylidene difluoride membrane (PVDF, Bio-Rad, cat. #1620177). Membranes were blocked for 5 min in Tris-buffered saline containing 0.1% Tween-20 (TBS-T) and 5% milk and incubated overnight in the same blocking buffer with one of the following primary antibodies: mouse monoclonal anti-LRRC8A (Santa Cruz, dilution 1:500) rabbit polyclonal anti-GLT-1 (Abcam, dilution 1:25,000). Blots were washed (3x) with TBS-T and probed with the species-matching secondary horseradish peroxidase-conjugated antibody: donkey anti-rabbit (GE Healthcare, dilution 1:10.000) or sheep anti-mouse IgG (GE Healthcare, 1:10,000) in TBS-T containing 5% milk for 2 h at room temperature. After a final wash, immunoreactivity was measured using ECL reagent (GE Healthcare, cat. #RPN2232) and visualized in a Bio-Rad ChemiDoc Imager (RRID:SCR_019684). For loading controls, membranes were re-probed with the HRP-conjugated primary anti-β-actin antibody for 20 min (Millipore-Sigma, 1:50,000), and the β-actin immunoreactivity was visualized.

### VRAC activity assay

VRAC activity was quantified by measuring the release of the non-metabolizable analog of glutamate, D-[^3^H]aspartate, as described and validated in our prior work (Abdullaev et al., 2006;Bowens et al., 2013;Schober et al., 2017). Primary astrocyte cultures from all relevant genotypes were plated on to poly-D-lysine treated 18×18 mm glass coverslips. Cells were loaded with 2 μCi/ml D-[2,3-^3^H]aspartate (Perkin Elmer, Waltham, MA, cat. #NET50100) in MEM +HIHS cell culture medium. Extracellular isotope was removed by washing in the chemically defined isoosmotic Basal medium containing (in mM) 135 NaCl, 3.8 KCl, 1.2 MgSO_4_, 1.3 CaCl_2_, 1.2 KH_2_PO_4_, 10 HEPES, and 10 D-glucose (pH 7.4, osmolarity 290 ±2mOsm). Coverslips were then transferred into a Leucite perfusion chamber with inlet/outlet ports. Cells were superfused with a either isoosmotic Basal medium or Hypoosmotic medium in which osmolarity was reduced to 200 ±2 mOsm (50-mM reduction in [NaCl]). One-minute perfusate fractions (~1.2 mL) were collected into scintillation vials via an automated fraction collector Spectra/Chrom CF-1 (Spectrum Chemical, New Brunswick, NJ, USA). At the end of sample collection, cell were lysed in solution of 2% sodium dodecyl sulfate (SDS) plus 8 mM EDTA. The [^3^H] content in individual samples was determined using a TriCarb 4910 TR scintillation counter (Perkin Elmer) after addition of scintillation liquid (Ecoscint, Atlanta Biologicals, Atlanta, GA, USA, cat. #LS-273). Outcomes were analyzed and reported as fractional release ratios relative to the total [^3^H] content for each time point, calculated using a custom Excel program.

### Middle cerebral artery occlusion surgery in mice

MCAo was performed as previously described with minor modifications (Balkaya et al., 2021). After a neck incision and dissection of left external carotid artery, a 6-0 silicon coated occlusion probe (Catalog No. 602356PK10, Doccol, Redland, CA, USA) was inserted and advanced into the internal carotid artery to block the blood flow at the origin of the middle cerebral artery. A fiberoptic cable (1mm diameter) was glued to the exposed skull (approximately 1.5 mm caudal, 4mm lateral of bregma) and the cerebral blood flow in the ischemic tissue was monitored in real time by laser-Doppler flowmetry (moorVMS-LDF1, Moor Instruments). Body temperature was monitored with a rectal probe in all animals and kept at 37 ±0.5°C. In this setting, brain temperature was 36.5 ±0.3°C as determined in a pilot group of mice using a probe placed in the brain. After 40-min occlusion, probe was withdrawn, and blood flow was restored. After ten minutes of reperfusion, neck incision was closed using surgical staples. Lidocaine hydrocloride jelly (2%, Akorn) was applied topically and Bupivacaine (0.25%, Auromedics) was injected subcutaneously as local analgesics post MCAo surgery. During recovery, additional analgesia was maintained by repeated local application of 2.5% Lidocaine/2.5% Prilocaine Cream (Actavis). We applied the following preregistered criteria to verify MCAo surgery success and minimize experimental variability. The inclusion criteria were (1) average reduction of cerebral blood flow by 75% or more during occlusion (equal or below 25% of the baseline measured with laser Doppler probe); (2) restoration and stabilization of blood flow to more than 65% of the original baseline within 10 minutes after filament withdrawal. The exclusion criteria were (1) insufficient reduction of blood flow during MCAo, (2) incomplete reperfusion (below targeted level), (3) animal death during recovery from anesthesia or within 1 hour after completion of MCAo. All included animals were allowed to survive for full 72 hours and euthanized after behavioral testing. Premature animal mortality between 1 and 72 hours was analyzed as a secondary outcome and reported in the paper.

### Brain lesion measurements

Ischemic lesion sizes were quantified at 24 or 72 h after initiation of ischemia using a triphenyl tetrazolium chloride (TTC) method (Bederson et al., 1986). TTC stains viable tissue in shades of red while the infarcted tissue remains unstained. Animals were deeply anesthetized via 5% isoflurane and decapitated. Brains were extracted and sliced rostro-caudally into serial 1-mm-thick slices using a Leica VT1200S vibratome (RRID: SCR_020243). Slices were submerged into 2% TTC solution in PBS and stained for 15-30 min at room temperature in a dark. Processed tissue specimens were fixed in 4% formaldehyde and stored at 4°C until scanned. Images were captured with a digital scanner and analyzed using ImageJ software (Rasband, 1997). The infarct volumes were corrected for brain edema and calculated by tracing healthy tissue in the ipsilateral hemispheres and subtracting it from contralateral hemisphere areas on all sections (Swanson et al., 1990).

### Evaluation of neurological deficits

While stroke lesion size can be used as a primary endpoint, there is a consensus in the field that behavioral outcomes are important as additional endpoints (Balkaya et al., 2018). Accordingly, we utilized tree different behavioral assessments (Neuroscore, Open Field test, and Pole test) to assess neurological recovery.

#### Neurological Score

MCAo animals were evaluated on a 5-point neurological score scale, immediately after recovery from anesthesia and daily until euthanasia. The scores were assigned as follows: 0, no neurological deficits; 1, failure to extend left forepaw; 2, circling to the left; 3, inability to carry bodyweight, falling to the left; 4, absence of spontaneous locomotion; 5, death.

#### Open Field test

Changes in spontaneous locomotor activity were tested in the open field, 72 hours post-surgery (Balkaya et al., 2013). Briefly, mice were released into the open field chamber (40 × 40 × 40 cm) and allowed to explore freely for 10 minutes. A video tracking software (AnyMaze) was used to track and analyze animals walking distance, time spent in the center vs the wall zones of the apparatus and the cumulative angle of rotation during locomotion. Only total travel distance is reported here.

#### Pole test

This test is a simple yet sensitive assay for evaluating motor function and coordination during the early post-stroke stages (Balkaya et al., 2013). Animals were trained to perform this task prior to MCAo. Ten rounds of training were generally sufficient for the test acclimation. Briefly, mice were placed on a 70-cm steel pole (1-cm thick) facing upwards and allowed to make a 180 degree turn and descend to the floor. For post-MCAo testing, the time to make a full turn (*tTurn*) and the total time to reach the floor (*tDescend*) was measured by analyzing video recordings of 5 successful trials and averaging their results. Mice that were too sick to perform (i.e., showing no spontaneous movement or repeatedly dropping from the pole) were assigned the highest scores among all the cohorts.

### Middle cerebral artery occlusion surgery and microdialysis in rats

Anesthesia was induced with 5% isoflurane in 30% O2/balance N2 and maintained through surgical procedures with 1.75-2.25% isoflurane under constant observation. To prevent excessive fluid secretion in the respiratory system and gastrointestinal tract, 0.4 mg/kg atropine sulfate was given via intramuscular injection. Animals were kept hydrated with hourly intraperitoneal (i.p.) injections of 1 ml physiological saline. Body and brain temperatures were maintained between 36.5°C to 37.5°C using a heating pad, and were measured with two probes placed rectally and in the temporalis muscle, respectively.

Two symmetrical bilateral microdialysis probes (CX-I series, 0.22×2 mm membrane, 50,000 kD molecular weight cutoff, Eicom Corporation, San Diego, CA) were lowered into the cortex through small burr holes, ~4 h before initiation of ischemia (see experimental design diagram in Figure 7). Probe placement was done using a stereotaxic frame (David Kopf Instruments, Tujunga, CA). The position of the probes was 2 mm anteroposterior, 5 mm lateral from bregma, and 2.6 mm down from dura. Artificial cerebrospinal fluid (aCSF) was perfused at a flow rate of 2 μl/min throughout the experiment. The composition of aCSF was as follows (in mM): 120 NaCl, 2.7 KCl, 1 MgSO4, 1.2 CaCl2, 25 NaHCO3, 0.05 ascorbic acid (pH=7.3). Microdialysis samples were collected every 20 minutes after a 1-1.5-h stabilization period using a CMA 470 refrigerated fraction collector (CMA Microdialysis, Holliston, MA). Cerebral blood flow was measured using a MoorLAB laser Doppler sensor mounted next to the microdialysis probe and acquisition module (Moor Instruments, Axminster, Devon, UK).

Middle cerebral artery occlusion (MCAo) was performed using an intraluminal suture technique as originally developed by Longa and colleagues (Longa et al., 1989) with modification detailed in our recent work (Dohare et al., 2014). Briefly, the right common carotid artery (CCA) and bifurcation of the external carotid and the internal carotid arteries (ECA and ICA) were exposed via incision on the neck. The ECA was coagulated. A standardized silicon-coated microfilament with a tip of 0.41 or 0.43 mm diameter (Doccol Corporation, Sharon, MA) was inserted via a stump of the ECA into the ICA up to 19-20 mm from the bifurcation to occlude the origin of MCA. The laser Doppler readings were used to ensure proper reduction of the blood flow rate to the levels that are expected in the ischemic penumbra. If blood flow was reduced by less than 80±1% (20% of the pre-ischemia values), animals were excluded from analysis on the basis of incomplete ischemia. After completion of ischemia and reperfusion in microdialysis experiments animals were euthanized with an overdose of sodium pentobarbital, brains were extracted and stained for early ischemic brain damage as described below.

### Analysis of microdialysate samples

Microdialysate samples were analyzed off-line. Amino acid levels in each sample were determined by a reverse-phase HPLC using an Agilent 1200 HPLC setup and Eclipse XDB-C18 column (both from Agilent Technologies, Santa Clara CA). Precolumn derivatization was performed with a freshly prepared mix of o-phthalaldehyde and 2-mercaptoethanol in 0.4 M sodium tetraborate buffer (pH = 9.5). The amino acid derivatives were eluted with solvent containing 30 mM NaH_2_PO_4_, 1% tetrahydrofuran, 30 mM sodium acetate, 0.05% sodium azide, and increasing concentration of HPLC grade methanol (10–30%). Fluorescence signal was measured using a programmable 1200 series fluorescence detector (Agilent). Amino acid standards (L-aspartate, L-glutamate, and taurine), were processed in the same fashion and used to identify amino acid peaks and calculate concentrations of individual amino acids in the samples.

### Sample size and statistics

Experimental outcomes in different genotypes have been statistically compared using one-way (effect of genotype) or two-way (effect of genotype and treatment) ANOVA with the Tukey or Sidak post-hoc tests for multiple comparisons, as appropriate. The effects of sex were not included in the main analysis but were additionally evaluated in a separate post-hoc assessment. For results involving multiple measurements, we used the repeated measures ANOVA or mixed model (Geisser-Greenhouse correction) with the Sidak’s post hoc test. Mortality values were evaluated using a Log-rank (Mantel-Cox) test. All statistical comparisons were performed using Prism 9.0 software (GraphPad Software Inc., La Jolla, CA). P values <0.05 were considered statistically significant.

The design of animal experiments was preplanned, including primary and secondary outcomes, and preregistered with the Open Science Framework [*https://osf.io/46npj* and *https://osf.io/3s7ab*]. The initial assumption was that nine animals per group would be sufficient to yield 80% power assuming that the maximal difference between groups will be at least 1.66 times the within group standard deviation (assumptions: alpha=0.05; parametric ANOVA; one factor, 4 levels; computed using Minitab software). This number (9/group) was further inflated to 15 mice/group based on the following assumptions. We expected ~10% surgical mortality, ~15% attrition based on stroke exclusion criteria, and additional ~15% mortality due to tamoxifen toxicity. As study progressed, we found that that some of the assumptions would have to be modified because we have higher-than-expected surgical attrition and stroke mortality. Furthermore, bLRRC8A KO mice have an inherited seizure phenotype, making it difficult to produce equal number of contemporary surgeries in each group. For these reasons we have completed stroke experiments when the histological testing of primary outcomes was ?9 mice per group.

## Abbreviations

CNS: central nervous system;
GFAP: glial fibrillary acidic protein;
GLT1: glutamate transporter 1;
LRRC8: leucine-reach repeat-containing family 8;
KO: gene knockout;
MCAo: middle cerebral artery occlusion;
4-OHT: 4-hydroxy-tamoxifen
VRAC: volume regulated anion channel

